# Tim-3 promotes viral infection by suppressing USP25-TRAF3-IRF7 signaling pathway

**DOI:** 10.1101/2024.08.19.608574

**Authors:** Lin Du, Jinjie Chen, Chunxiao Du, Junrui Chen, Chaoxiang Wang, Bing Bao, Zhonglin Lv, Chen Xing, Meng Liang, Lanying Wang, Shun Xie, Yuxiang Li, Zhiding Wang, Ge Li, Jun Zhang, Gencheng Han

**Author notes:** These authors contributed equally to this work (Lin Du, Jinjie Chen).

## Abstract

Tim-3, an immune checkpoint inhibitor, plays key roles in maintaining immune homeostasis and is involved in viral evasion. However, the precise role of Tim-3 in viral infection remains to be determined. USP25 is one of the deubiquitinating enzymes that serves to initiate antiviral immunity by deubiquitinating TRAF3 and triggering the anti-viral signaling pathway. Here we found that Tim-3 specific knockout in myeloid cells leads to enhanced anti-viral immunity in mice with VSV encephalitis by increasing type I interferon response. Mechanistically, Tim-3 inhibits the expression of USP25 via STAT1, interacts with USP25 but does not regulate its post-translational modification, and as a result, Tim-3 inhibits USP25 mediated deubiquitination of TRAF3, promotes k48-linked ubiquitination and degradation of TRAF3, and then inhibits the phosphorylation of IRF7, finally down-regulates the interferon response. These findings provide new insights into the function of Tim-3 in antiviral immunity and its related clinical significance.

## Introduction

T cell immunoglobulin and mucin domain 3 (Tim-3) is widely expressed on many immune cells including T cells and macrophages^[1–4]^. Establishment of Tim-3 as an exhaustion marker in immune cells of both tumors and infectious diseases makes Tim-3 an attractive target for immunotherapy similar to PD-1 and CTLA-4. Recently, a report showed that increased Tim-3 expression on immune cells in patients with coronavirus disease (COVID-19) is associated with an exhaustion phenotype^[5]^. Compared to the relatively clear mechanisms by which PD-1 induces tolerance in T cells, very little is known about Tim-3 signaling in immune cells especially in innate immune cells. We previously demonstrated that Tim-3 inhibits NF-kB activation in macrophages in response to LPS stimulation^[6]^ and that Tim-3 promotes M2 macrophage polarization by inhibiting the activity of STAT1^[7]^. We also found that Tim-3 controlled the response of macrophages by enhancing the ubiquitination and degradation of MHC-I, and Tim-3 negatively regulated the macrophage-mediated antigen presentation^[8]^. These data suggest that Tim-3 play a key role in regulating the innate immune response. Tim-3 does not have an inhibitory motif within its tail, and the mechanism by which Tim-3 mediates inhibitory signaling remains largely unclear^[9; 10]^. Potentiating anti-infection immunity by inducing innate immune responses is a promising area of infection therapy. However, in case of viral infection, little is known about Tim-3 signaling in innate immune cells.

Deubiquitinating enzymes (DUBs) are capable of recognizing and cleaving peptide bonds between ubiquitin and substrate or within the ubiquitin chain, which have multiple biological activity^[11]^. Recently, studies have found that ubiquitin specific proteases family molecule 25 (USP25) is involved in the pathogenesis of various diseases including infectious diseases, neurodegenerative disorders, and cancer^[12–15]^. Lin et al. found that compared with wild-type mice, USP25-deficient mice were more susceptible to infection with herpes simplex virus 1 or influenza A virus subtype H5N1^[16]^. Studies showed that USP25 promotes the antiviral immune response by cleaving ubiquitin moieties from critical signaling proteins of the type I IFN signaling pathway such as RIG-I, TRAF3, and TRAF6^[17]^. USP25 tends to play an essential role in infection disease, but many of the mechanisms are not fully detailed.

Innate immunity is the first defense line against viral infection. A variety of effector cells in the innate immune system can quickly recognize pathogens and produce type I interferon and other cytokines to inhibit virus replication and transmission^[18; 19]^. Interferon regulatory factor 7 (IRF7), which is the signaling cascade downstream of TRAF3, plays ciritical roles in anti-viral innate immunity by regulating the response of type I interferon, interferon stimulated gene (ISG) and other pro-inflammatory cytokines. The signaling activity of IRF7 is regulated by multiple mechanisms. Investigation on the mechanisms by which IRF7 is regulated among different physiopathological conditions will provide much need information for anti-viral innate immunity.

Here we identified a new mechanism by which Tim-3 promotes immune evasion, that is, by suppressing USP25 expression, Tim-3 inhibits USP25 mediated deubiquitinating of TRAF3 and suppresses the TRAF3-IRF7-Type I interferon pathway. We thus identified a new signaling pathway of Tim-3 mediated immune tolerance with clinical significance.

## Results

### Specific knock out of Tim-3 in myeloid cells mice resulted in increased survival rate in mice with VSV encephalitis

To investigate the roles of Tim-3 in anti-viral innate immunity, Tim-3 myeloid specific knockout mice (Tim-3-CKO) and wild type mice (WT) were induced with encephalitis via Vesicular Stomatitis Virus (VSV) and then the survival rate and immune response were checked. We found Tim-3 myeloid specific knockout leads to increased survival rate following VSV encephalitis (Fig.1A). We then examined the effects of Tim-3 myeloid knock out on tissue damage in mice with VSV encephalitis. Fig.1B showed that compare to wild type mice, Tim-3-CKO mice with VSV encephalitis showed markedly decreased meningeal injury as marked by the decreased perivascular cuffing. When the viral replication in mice with or without Tim-3 knockout were examined, Fig.1C&D showed that Tim-3-CKO lead to decreased viral replication both in the brain and PBMC in mice with VSV encephalitis. Finally, we examined the effects of Tim-3-CKO on the neuroethology in mice with VSV encephalitis using open-field test (OFT). The results showed that the line crossing was significantly higher in Tim-3-CKO mice than that in wild type mice with VSV encephalitis (Fig.1F). In addition, Tim-3 myeloid specific knockout lead to much higher average speed and total distance in the open field test (Fig.1G&H). These results indicated that Tim-3 myeloid specific knock out lead to suppressed viral replication and attenuated symptoms of encephalitis.

**Figure 1.**
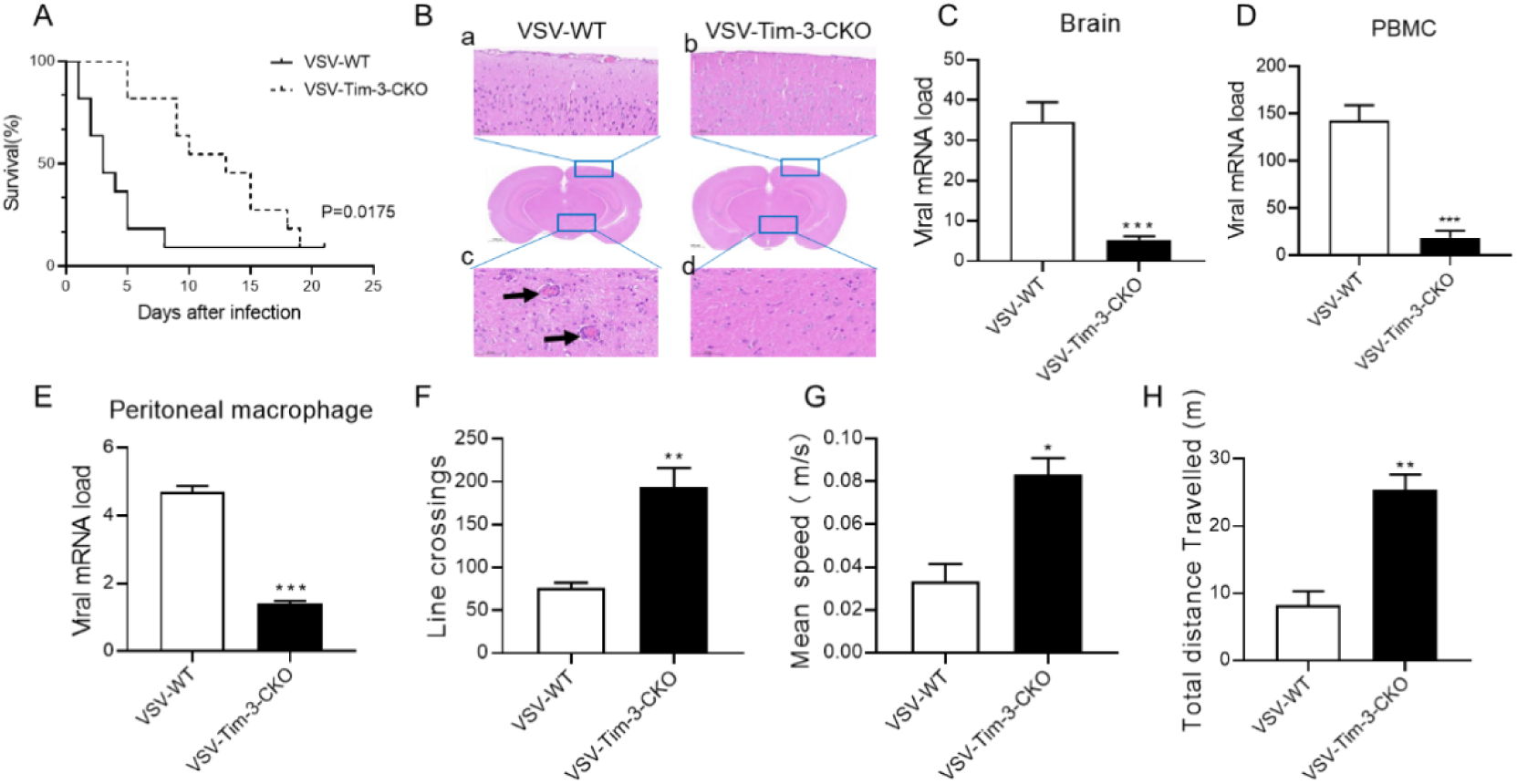
Specific knock out of Tim-3 in myeloid cells protects mice from VSV encephalitis. (A) Wild type mouse and Tim-3-CKO mice were intracranially injected with VSV (1X10^6^ pfu/g).Survival rate was analyzed (n=10). (B) Mice were infected and treated as elaborated in (A). Brain tissues were collected on day 5 post-infection and stain with hematoxylin and eosin. Arrow in a) shows the damaged meningitis which is absent in b). arrow in c) shows perivascular lymphocyte cuffing which is a well-known indicator of inflammation in the brain. (C-E) Brain, PBMC and peritoneal macrophage were collected on day 5 post-infection and VSV replication were analyzed by RT-PCR(n=10, Mean ± SD). (F-H) Mice were infected and treated as elaborated in (A). At day 5, mice underwent the open-field test, and their spontaneous locomotor activity was evaluated by measuring the line crossing, mean speed and the distance traveled in a defined area for 10 min (n=6~8, Mean ± SD). The results shown are representative of three independent experiments. *p<0.05, **p<0.01, ***p<0.001.

### Tim-3 myeloid specific knock out increased the production of type I interferons in mice with VSV encephalitis

It is known that interferons signaling play critical roles in anti-viral innate immunity. To find the mechanisms by which Tim-3 signaling promotes VSV replication, the brain tissues and PBMC samples were collected from WT and Tim-3-CKO mice with VSV encephalitis and the expression of the IFN-α4 and IFN-β were detected by Real-time PCR (RT-PCR). The data in Fig.2 showed that the Tim-3 myeloid specific knockout lead to increased expression of IFN-α4 and IFN-β in mice following VSV infection, suggesting that Tim-3 might exacerbate VSV encephalitis via suppressing the interferon signaling pathway.

**Figure 2.**
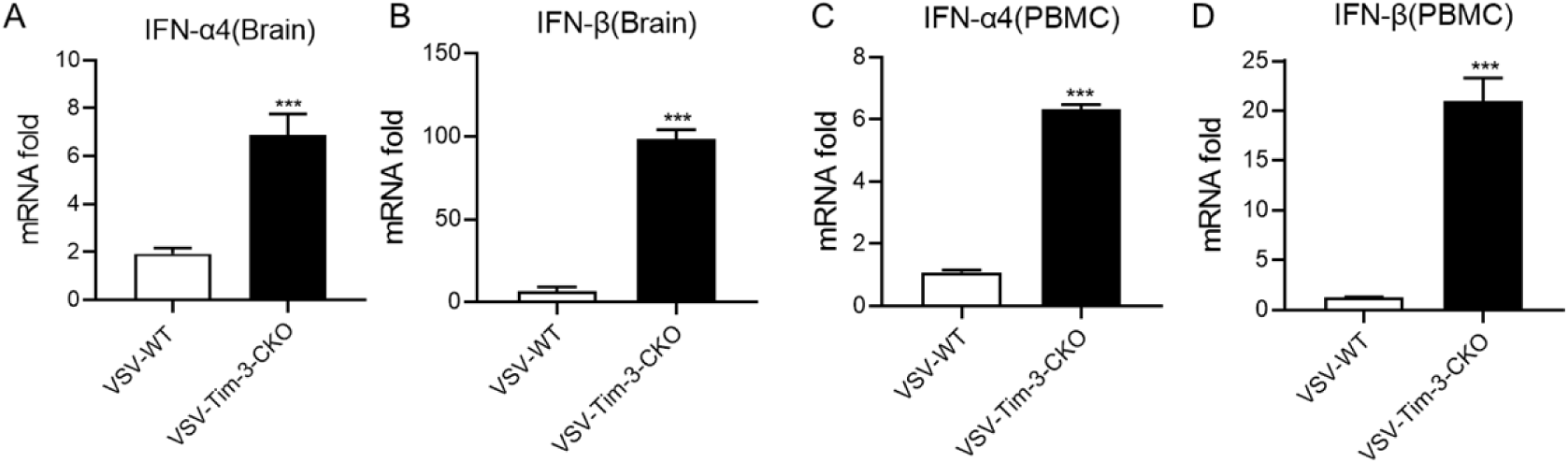
Tim-3 myeloid specific knock out mice showed increased the production of type I interferon (IFNs) in mice with VSV encephalitis. Mice were infected and treated as elaborated in Fig.1. At day 5 mice were sacrificed and brain tissue and PBMC were collected. (A&B) Expression of type I interferon factors of IFN-α4 and IFN-β in the brain of mice. (C&D) Expression of type I interferon factors of IFN-α4 and IFN-β in the PBMC of mice. n=10 for each group. Data are presented as mean± SEM. The results shown are representative of three independent experiments. ***p<0.001.

### Tim-3 signaling inhibits type I IFN expression through USP25

USP25 is a deubiquitinating enzyme which has been reported to play an important role in antiviral immunity. As the analysis on GEO dataset (GSE50011) for 86 infected individual samples showed that there is an inverse correlation (r = −0.58868, p<0.0001) between the expression of Tim-3 and USP25 in CD14^+^ monocytes/macrophages (Fig. 3A). We hypothesized that Tim-3 might regulate the USP25-type I IFN signaling pathway. To test this hypothesis, we silenced USP25 in macrophages and examined type I IFN expression with or without Tim-3 blockade. When USP25 was silenced (Fig. 3B), Tim-3 blockade did not increase IFN-α4 and IFN-β expression again (Fig. 3C&D). These results indicate that Tim-3 may inhibit type I IFN expression through USP25.

**Figure 3.**
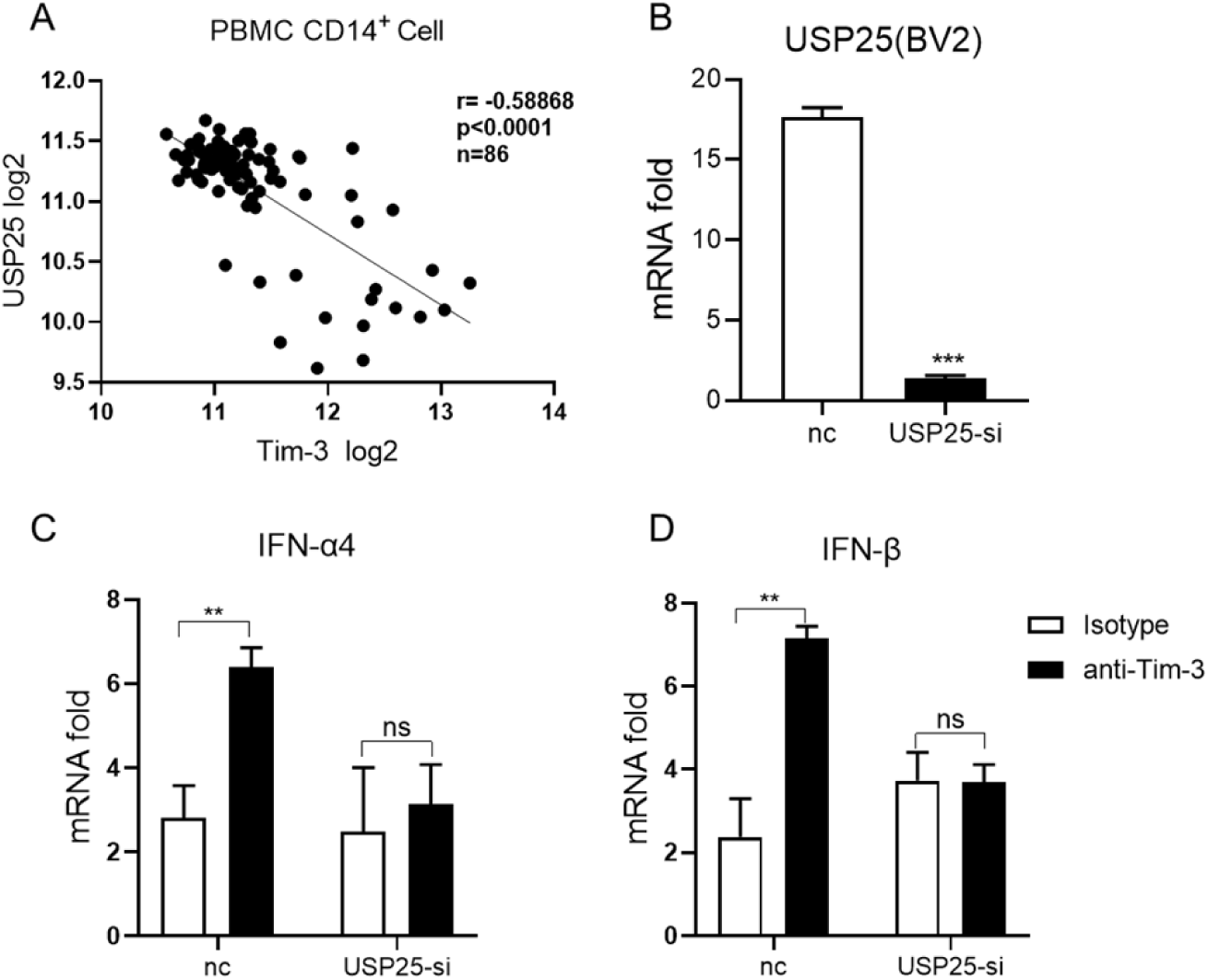
Tim-3 inhibits USP25 expression both in vivo and in vitro in response to VSV infection. (A) The expression data of Tim-3 and USP25 in CD14^+^ monocytes/macrophages from 86 infected individual samples were collected from the GEO dataset (GSE50011), and their correlation was analyzed. (B) BV2 cells were transfected with control siRNA (nc) or USP25 siRNA (USP25-si) for 24h. Then the expression of USP25 was analyzed using RT-PCR. (C&D) The expression of IFN was analyzed by RT-PCR. Data are presented as mean± SD. The results shown in B-D are representative of three independent experiments. **p<0.01, ***p<0.001.

### Tim-3 inhibits USP25 expression both in vivo and in vitro in response to VSV infection

The above GEO dataset showing that Tim-3 negatively correlated with USP25.The effects of Tim-3 on USP25 expression was further confirmed both in VSV infected mice in vivo and in VSV infected cells in vitro. The data in Fig.4A-C showed that Tim-3 myeloid specific knockout lead to increased USP25 expression in mice with VSV encephalitis. In addition, when Tim-3 is overexpressed in HEK293T cells or is silenced in macrophage cell line RAW264.7 cells or silenced in microglia cell line BV-2 cells, the data in Fig.4D-F showed that, in response to VSV infection, Tim-3 suppresses USP25 expression both at protein and mRNA level.

**Figure 4.**
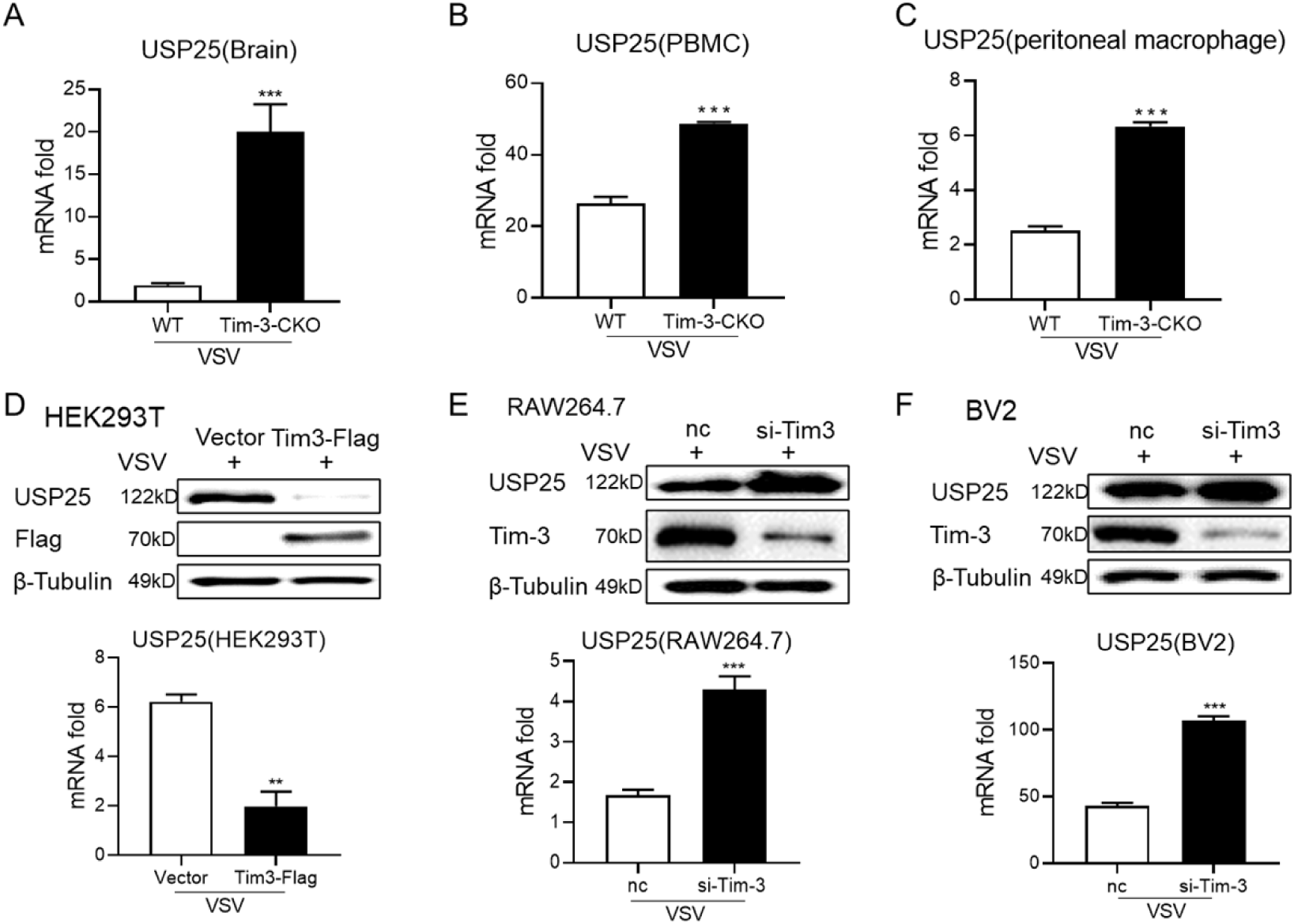
Tim-3 inhibits USP25 expression both in vivo and in vitro in response to VSV infection. Mice were infected and treated as elaborated in Fig.1. (A-C) Brain, PBMC and peritoneal macrophage were collected on day 5 post-infection, the expression of USP25 were analyzed by RT-PCR. (D) HEK293T cells were transferred with Tim-3-Flag plasmid. Following VSV infection for 12h USP25 protein (upper panel) and gene levels (lower panel) were detected by western blot and RT-PCR at 48 hours following transfection. (E&F) Tim-3 was knockdown by shRNA in RAW264.7 cells (E) or in BV2 cells (F). These cells were then challenged with VSV for 12 hours. The effects of Tim-3 on USP25 expression were examined by western blot (upper panel) and RT-PCR (lower panel). The results shown are representative of three independent experiments. **P<0.01, ***P<0.001.

### Tim-3 signaling inhibits USP25 expression through STAT1

The mechanisms by which Tim-3 suppresses USP25 expression were then investigated. As we previously found that translational factor STAT1 might act as the intracellular signaling adaptor of Tim-3^[7]^, we then tested whether Tim-3 suppresses USP25 via STAT1. We found that transfection of STAT1 in HEK293T cells increased the expression of USP25 at both protein and mRNA level (Fig.5A). However, co-transfection of Tim-3 with STAT1 reverses STAT1 increased USP25 expression (Fig.5B). Luciferase reporter assay showed that co-transfection of Tim-3 with STAT1 reverses transfection of STAT1 alone induced up-regulation of USP25 (Fig.5C). Furthermore, we found that co-transfection of Tim-3 with STAT1 inhibitor fludarabine reverses Tim-3 mediated suppression on USP25 in HEK293-T cells (Fig.5D). These data demonstrated that Tim-3 suppresses USP25 expression via STAT1.

**Figure 5.**
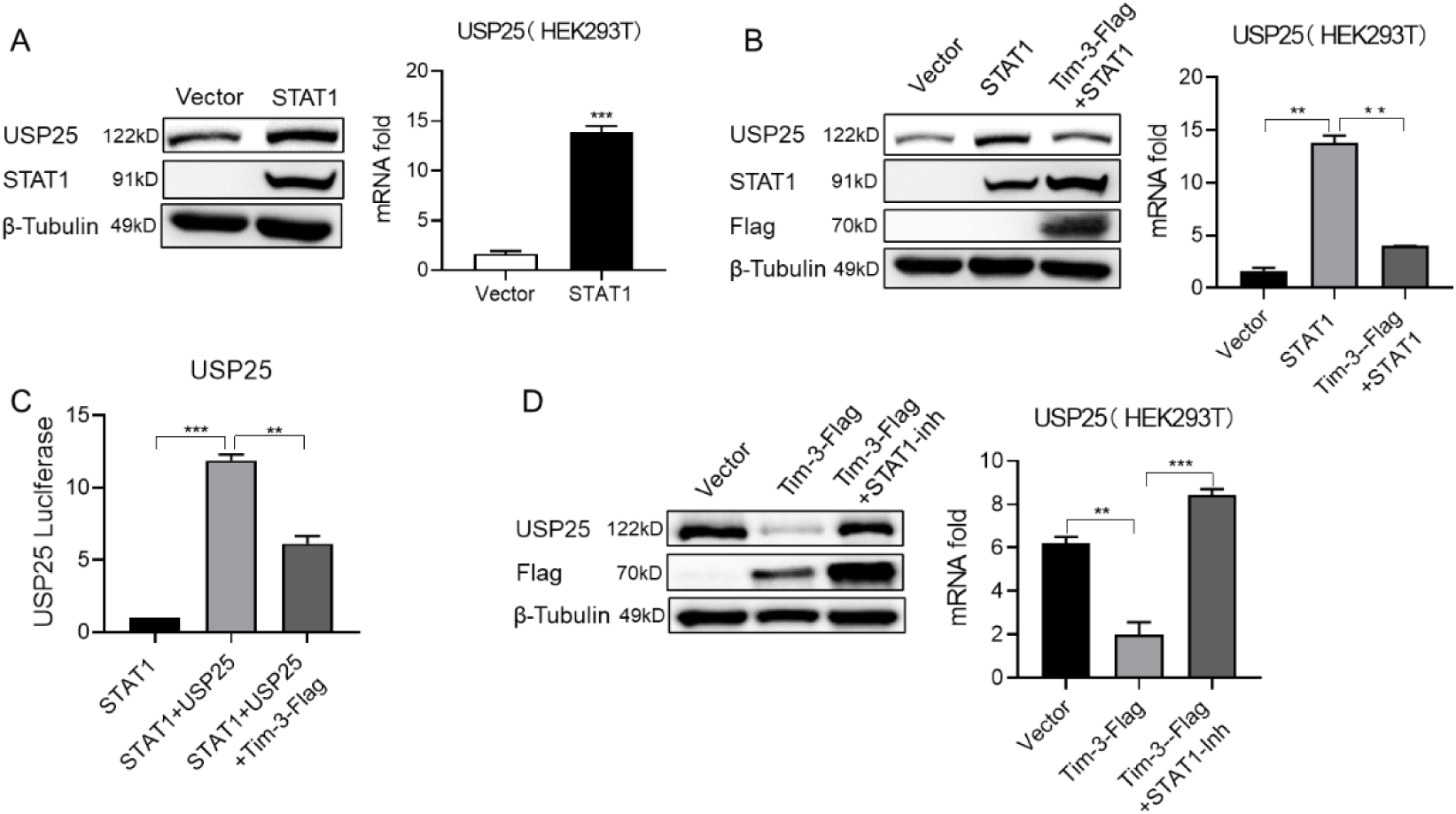
Tim-3 signaling inhibits USP25 expression through STAT1 in macrophages. **(A)** HEK-293T cells were transferred into STAT1 plasmid. Then USP25 expression was detected by western blot (left panel) and RT-PCR (right panel) at 48 hours following transfection. (B) HEK-293T cells were transferred with STAT1 plasmid, with or without Tim-3 plasmid and lysed, and the protein levels (left panel) and gene levels (right panel) of USP25 were analyzed at 48 hours following transfection. (C) HEK-293T cells were transferred into pGL3-USP25 reporter plasmid and STAT1 plasmid, with or without Tim-3 plasmid for 48 hours following transfection. Dual-fluorescence was analyzed. (D) HEK-293T cells were transferred into Tim-3-flag plasmid, and in the presence of the STAT1 inhibitor fludarabine (10 μm), and USP25 protein levels (left panel) and the mRNA levels (right panel) were analyzed at 48 hours following transfection. The results shown are representative of three independent experiments **P<0.01, ***P<0.001.

### Tim-3 inhibits USP25 mediated deubiquitination of TRAF3 and suppresses the phosphorylation of IRF7

The above data showed that Tim-3 inhibits the transcription of USP25 via STAT1. We then examined whether or how Tim-3 regulates signaling cascade downstream of USP25. First, we investigated whether Tim-3 interacts with and/or is involved in the post-translational modification of USP25. HEK293T cells were co-transfected with USP25 and Tim-3, then immunoprecipitation was conducted targeting either Tim-3 (Fig.S1A) or USP25 (Fig. S1B). All of them confirmed the direct interaction between Tim-3 and USP25. However,, we found that Tim-3 is not involved in the post-translational regulation of USP25, as the data in Fig. S1C show that transfection of Tim-3 did not decreased the half-life of endogenous USP protein within 6 h in the presence of cycloheximide (CHX), a ribosome inhibitor that blocks protein synthesis in ribosomes. Collectively, these results suggest that Tim-3 interact with USP25 but have no affect with its post-translation modification.

We then examined the effects of Tim-3 on USP25 mediated deubiquitination of TRAF3. As shown in Fig.6A, when USP25 and TRAF3 were co-transfected into HEK293T cells, the ubiquitination of TRAF3 was significantly decreased. While when Tim-3 is co-transfected with USP25 and TRAF3, Tim-3 inhibits USP25 mediated de-ubiquitination of TRAF3. As K48-linked ubiquitination plays a critical role in the function of TRAF3, we then examined whether Tim-3 inhibits USP25 mediated deubiquitination of TRAF3 following K48-Ub transfection. The data in Fig.6B showed that USP25 significantly decreases the K48-linked ubiquitination of TRAF3, which can be reversed when Tim-3 is co-transfected. These data suggested that Tim-3 suppresses the USP25-TRAF3 signaling pathway.

**Figure 6.**
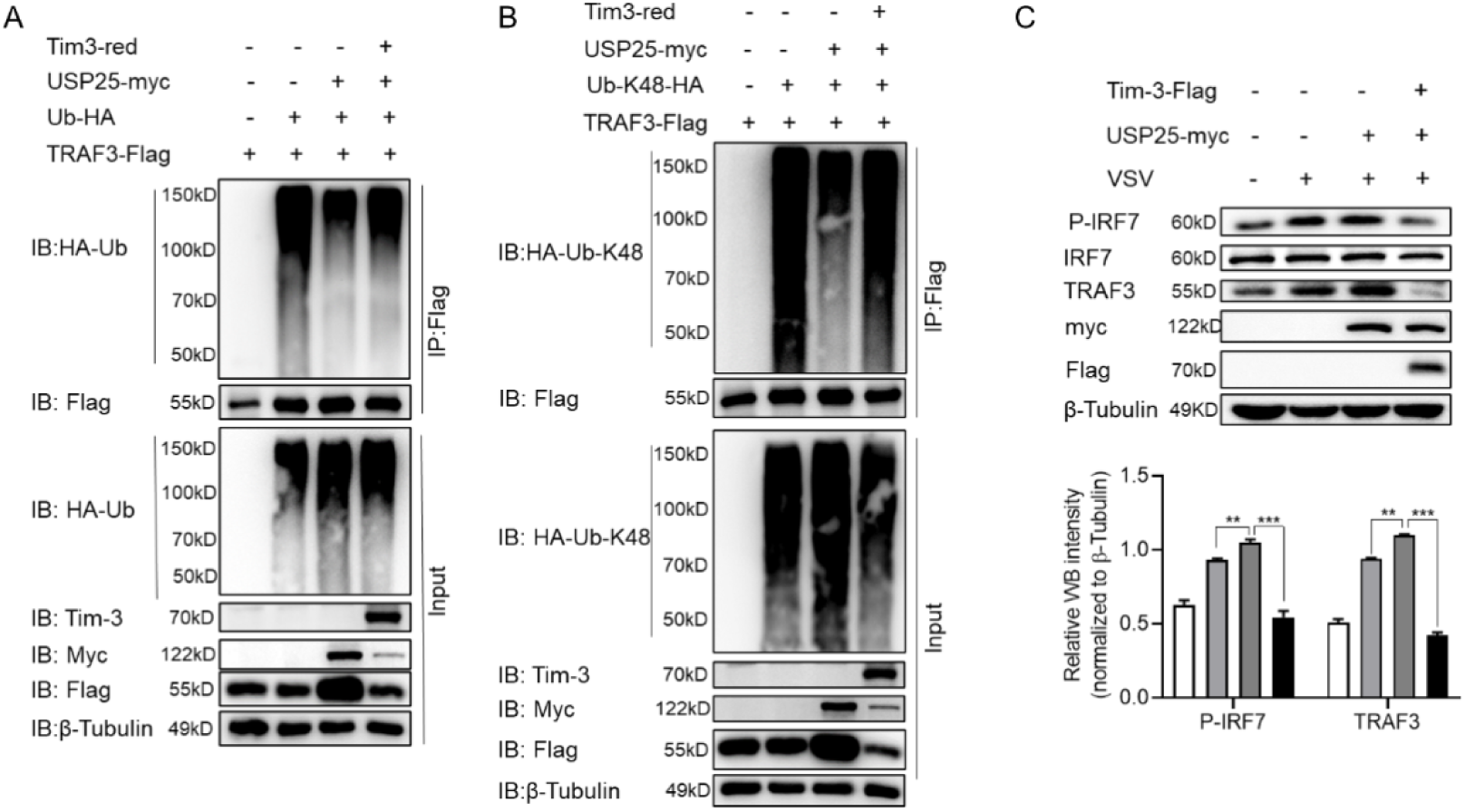
Tim-3 controls the ubiquitination of TRAF3 and the phosphorylation of IRF7 via USP25 in macrophage. (A) Plasmids encoding HA-tag-ubiquitin, USP25-myc, TRAF3-Flag and Tim-3-red were transfected into HEK293T cells. Cells were treated with MG132 (20 µg/ml) for 6 hours before harvesting protein lysates, followed by immunoprecipitation with antibody to Flag and then Western Blot analysis of ub-linked TRAF3 ubiquitination with antibody to HA. (B) Plasmids encoding HA-tag-ubiquitin-K48-lined ubiquitin, USP25-myc, TRAF3-Flag and Tim-3-red were transfected into HEK293T cells. Cells were treated with MG132 (20 µg/ml) for 6 hours before harvesting protein lysates, followed by immunoprecipitation with antibody to Flag and then Western Blot analysis of K48-linked TRAF3 ubiquitination with antibody to HA. (C) HEK-293T cells were transferred into Tim-3-flag plasmid and USP25-myc plasmid, with or without VSV infected for 12 hours, the phosphorylation of IRF7 and expression of TRAF3 were analysis by western blot cohort (upper panel) along with quantification of results (lower panel). The results shown are representative of three independent experiments. **P<0.01, ***P<0.001.

As an important signaling cascade down-stream of TRAF3, IRF7 plays critical role in interferon response. To test whether Tim-3 signaling down regulates interferon response via USP25-TRAF3-IRF7 pathway, we co-transfected HEK293T cells with Tim-3 and USP25 in the presence or absence of VSV. The data in Fig.6 C showed that co-transfection of Tim-3 with USP25 reverses USP25 increased TRAF3 expression and reverses USP25 increased IRF7 phosphorylation. As IRF7 is a critical signaling cascade regulating interferon response, these data partially explained the mechanisms by which Tim-3 inhibits interferon response in VSV encephalitis mice in vivo.

## Discussion

Here we found a new mechanism by which Tim-3 promotes immune tolerance and mediates virus evasion. First, we found that Tim-3 inhibits the transcription of USP25, one of the deubiquitinating enzymes which has been reported to enhance anti-viral immune response. As a result, Tim-3 inhibits USP25 mediated deubiquitination of TRAF3, promote k48-linked ubiquitination of TRAF3 and then inhibits the phosphorylation of IRF7, the signaling cascade down-stream of TRAF3. IRF7 play critical roles in type I interferon response. Our results explained why Tim-3 signaling inhibits interferon response and mediates infection tolerance in mice with VSV encephalitis. These findings provide insights into the function of Tim-3 in antiviral innate immunity and may have clinical significance.

USP25 is involved in the processes by which toll-like receptors (TLR) monitor and recognize a variety of different disease-associated molecular patterns and provide the first barrier of the body against infectious diseases. Upon infection with an RNA or DNA virus, USP25 associates with TRAF3 and TRAF6 and protects TRAF3 and TRAF6 from virus-induced proteasome-dependent or independent degradation^[16]^. For example, upon stimulation of TLR4 by LPS, USP25 was shown to specifically reverse the Lys48-linked ubiquitination of TRAF3 mediated by the E3 ubiquitin ligase inhibitor of apoptosis (Ciap2)^[20]^. Li et al. found that USP25 contains conserved Cys178 residues, which are responsible for the deubiquitination activity of USP25^[21; 22]^. USP25 protected TRAF3 and TRAF6 from degradation and thus positively regulated virus-induced production of type I IFNs and proinflammatory cytokines^[20]^. In this study, we found that Tim-3 inhibits USP25 expression and down-regulates type I interferon expression in mice with virus infection. Mechanistically, Tim-3 inhibits USP25 mediated deubiquitination of TRAF3 and then suppresses the phosphorylation of IRF7^[17; 23]^. However, although our data demonstrate the interaction between Tim-3 and USP25, Tim-3 is not involved in the post-translation regulation of USP25 (Fig.S1). Interestingly, the crystal structure of USP25 shows an unexpected combination in which a Hom tetramer is formed by the union of two dimers. This quaternary structure leads to a unique autoinhibitory mechanism in which the autoinhibitory motif of USP25 directly occupies a large portion of the ubiquitin-binding surface, ultimately leading to the inhibition of its enzymatic activity^[24; 25]^. However, whether Tim-3 affects the conformation of USP25 has not been carried out in this paper and will be investigated in the future.

Tim-3 is an immune checkpoint inhibitor that was initially identified on activated T cells, including Th1 and Th17 cells. Recently, checkpoint inhibitors PD-1 and CTLA-4 have been used as therapeutic targets for immune disorders including tumors and autoimmune disease. Tim-3 is considered as the next generation therapeutic target. We focused on the role of Tim-3 in maintain the homeostasis of innate immunity. So far, we have reported several pathways for Tim-3 mediated innate immune tolerance. For example, we found that Tim-3 promotes the ubiquitination and degradation of NF90 and inhibits NF90-SG-mediated antiviral immunity^[26]^. In addition, we also found that Tim-3 can inhibit the expression of RIG-I, which is a cytoplasmic sensor that triggers the induction of type I IFN production, leading to viral immune evasion^[27]^. These findings confirm the significant impact of Tim-3 on the antiviral immune response during infection. It is well-established that Tim-3 lacks an inhibitory motif within its tail^[28]^, and the precise mechanism by which Tim-3 mediates inhibitory signaling remains unclear. Our study here reveals that Tim-3 suppress the expression of USP25, a crucial regulator of innate immune responses, and then inhibited the expression of type I interferon, finally leading to viral immune escape. The biology of Tim-3 is intricate, involving non-canonical signaling, widespread expression across various types of immune cells, and interaction with multiple ligands. Although much remain to be learned about the roles of Tim-3 in immune homeostasis, here we found a new mechanism by which Tim-3 induces infection tolerance.

It is well known that interferon responses play critical roles in antiviral immunity. Research indicates type I IFN signaling activation relay on TRAF3 ubiquitin then facilitates phosphorylation of IFN regulatory factor (IRF)3/7 and translocation into nucleus^[29; 30]^. TRAF3 selectively activates the production of proinflammatory cytokines and type I IFNs through K48- or K63-linked ubiquitination events, respectively^[31]^. Many viruses developed ways to escape immune attack by suppressing interferon response. For example, the non-structural protein 1 of influenza A viruses inhibits the effects of IFN mainly via blocking the expression of IFNs to facilitate influenza virus evasion^[32; 33]^. Here we found a new mechanism by which virus suppresses interferon response, that is Virus up-regulates the expression of Tim-3, the increased Tim-3 in turn suppress USP25-TRAF3-IRF7-Type I interferon signaling and finally leading to viral immune escape.

In summary, we identified a novel mechanism by which Tim-3 induces immune tolerance and promotes virus evasion, Tim-3 interacts with USP25, inhibits USP25 expression through STAT1, inhibits IRF7 phosphorylation and promotes TRAF3 ubiquitination, and subsequently inhibits type I IFN production and antiviral activity. Our findings provide insight into the underlying mechanism of immune response against viral encephalitis and may have potential therapeutic applications.

**Figure 7.**
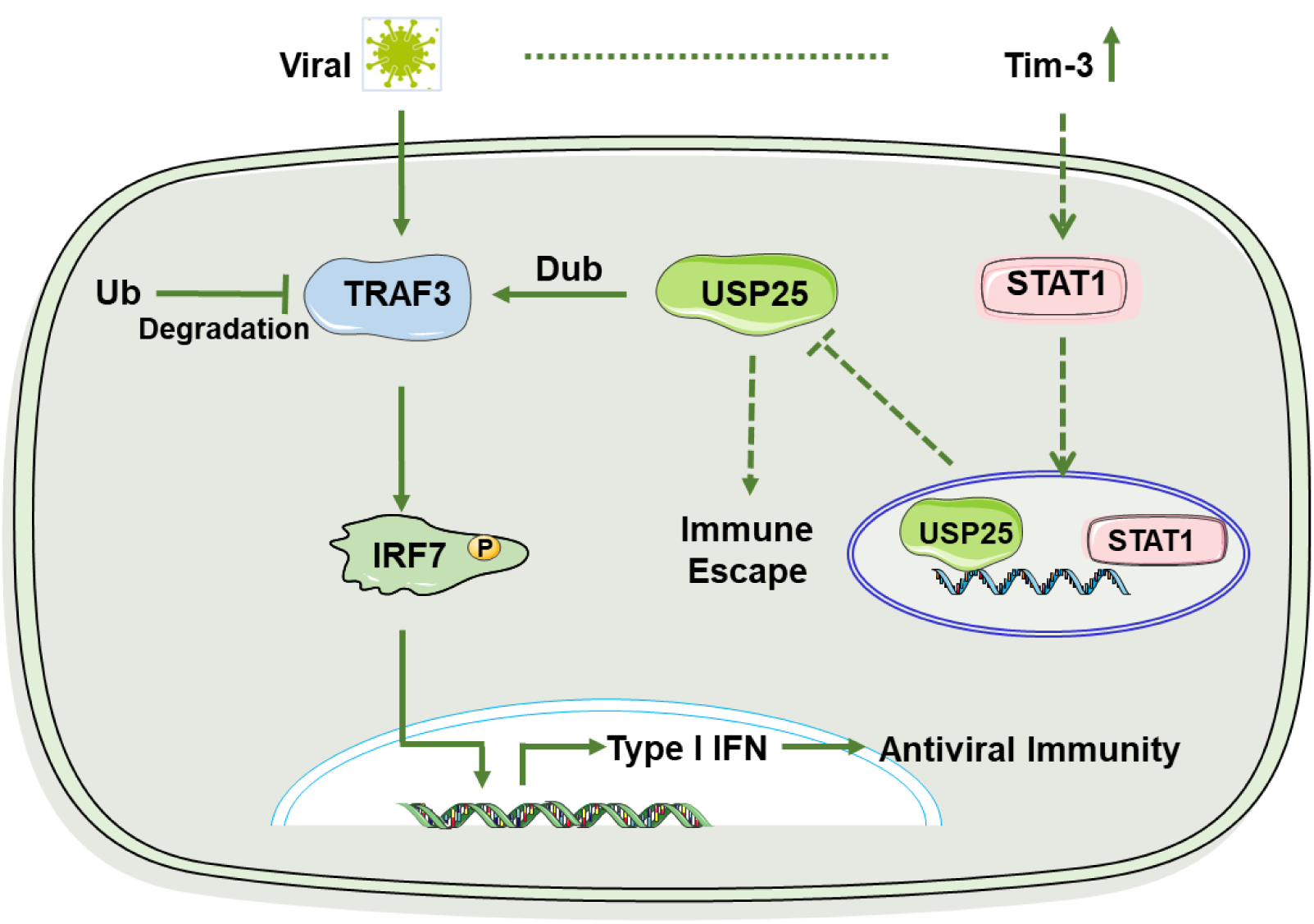
A schematic diagram illustrating how Tim-3 inhibits USP25-TRAF3-IRF7 pathway during viral infection. Upon vesicular stomatitis virus (VSV) infection, the increased Tim-3 inhibits USP25 expression through STAT1. As a result, Tim-3 inhibits USP25 mediated deubiquitination of TRAF3 and then inhibits the phosphorylation of IRF7, finally down-regulates the antiviral interferon response.

## Materials and methods

### Animals

Male C57BL/6J (6-8 weeks old) were purchased from Spaifer and Tim-3 knockout mice were purchased from Segen’s Biosciences (Guangzhou, China). All mice were bred and maintained under specific pathogen-free conditions, and the experimental was approved according to the protocol by the Ethics Committee of Animal Experiments of the Institute of Military Cognitive and Brain Sciences (permit number: IACUC-DWZX-2023-P526). All efforts were made to minimize animal suffering.

### In vivo Experimental VSV Infection

VSV was a gift from Prof. Minghong Jiang at the institute of Basic Medicine, Chinese Academy of Medical Sciences. Tim-3-CKO mice and WT mice were given intracranial injection with VSV (1 × 10^6^ pfu/g). The procedure was described in the previous study^[8]^.

### Open field test (OFT)

The open field chamber was made of transparent plastic (50 cm × 50 cm) and a 25 cm × 25 cm center square was color marked. Individual mice were placed in the center of the chamber and their behavior was monitored for 5 min with an overhead video-tracking system (ANYmaze, Ugo Basile, Varese, Italy). The time mice spent in the center area, the total distance they traveled and their velocity were monitored throughout the experiment.

### Cell culture and Transfection

BV2, RAW264.7, and HEK293T cells were from ATCC, si-Tim-3-RAW264.7 cells and si-Tim-3-BV2 cells (stably knocked out Tim-3) come from our laboratory. Mouse peritoneal macrophages were collected as previously described. All cells were maintained in Dulbecco’s Modified Eagle’s Medium (DMEM; Gibco, USA) supplemented with 10% fetal bovine serum, streptomycin (50μg/ml), and erythromycin (5μg/ml) were cultured in an incubator (37°C, 5% CO2).

For transient expression, the vector (pcDNA3.1) alone, Tim-3, STAT1, and USP25 plasmids were transfected into HEK293T cells alone or in combination and cultured for 48h after transfection. The plasmid was previously stored in our laboratory.

### H&E Staining

Brain tissues were excised from mice sacrificed on day 6 after infection, fixed with 4% paraformaldehyde, embedded in paraffin, and sectioned. The injury status of Brain in mice was assessed by Hematoxylin-eosin staining.

### Quantitative reverse transcription-PCR (RT-PCR)

The expression of VSV load, IFNα4, IFNβ and USP25 was analyzed by three-step reverse transcriptase quantitative-polymerase chain reaction (RT-PCR). Total RNA was extracted using TRlzol reagent (CWBIO) and reversed using the TransGen Biotech Reverse Transcription Kit in a 20 ml reaction volume (50°C, 5 min; 85°C, 15 s) according to the manufacturer’s instructions. Then cDNA was amplified using 2X Realstar Green Fast Mixture (Genstar) and a LightCycler 480 polymerase chain reaction system (Roche). All trials are performed in replicate reaction mixtures in 96-well plates. The relative expression of the gene of interest was determined by the 2-DDCT method of 18S ribosome, with 18S used as a control. The primer sequences are list in Table 1.

**Table 1.**
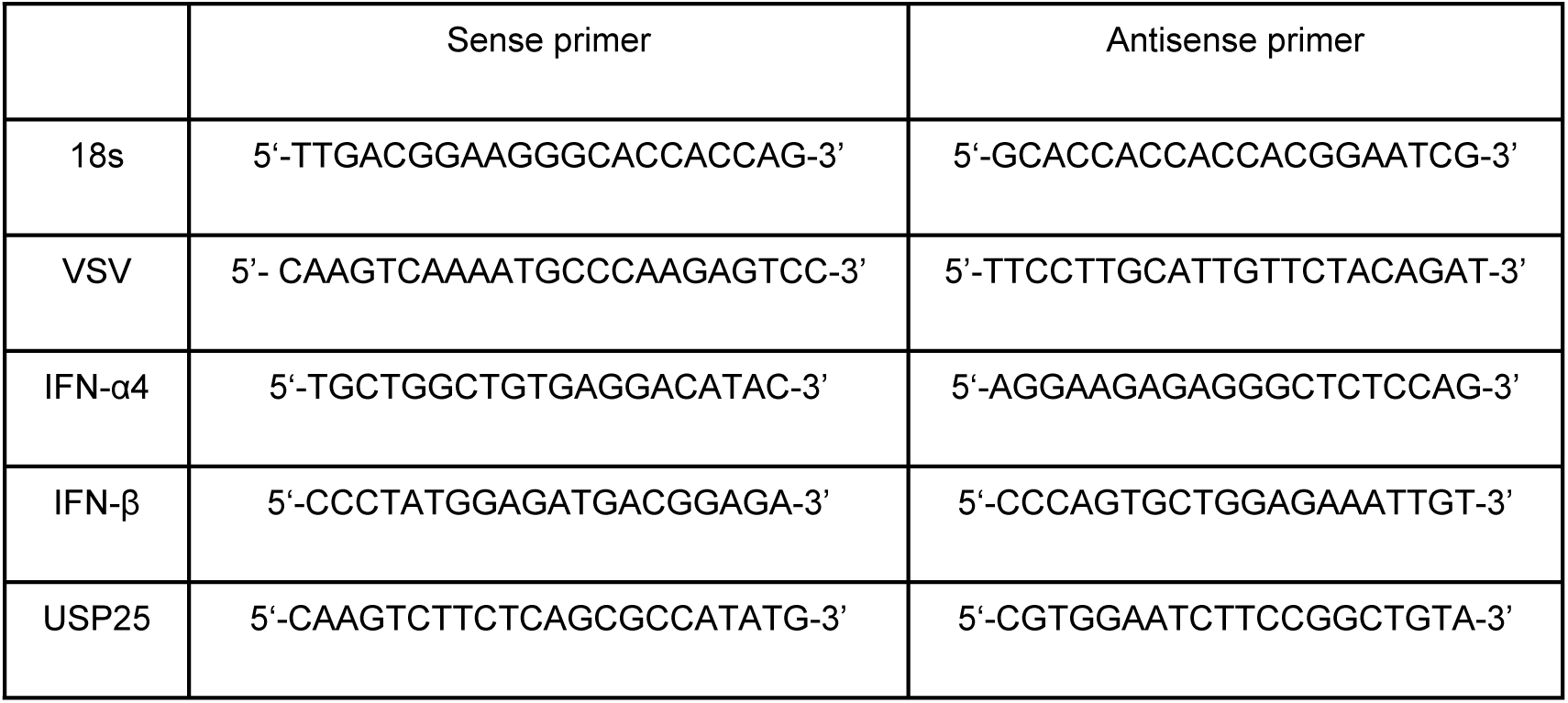
Sequences of the primers used for PCR.

### Western Blotting

Cells were harvested and lysed in lysis buffer supplemented with protease and phosphatase inhibitors, and collected from supernatant by centrifugation, followed by addition of loading buffer, boiling for 10 min and storage at −20°C. The protein concentration of the samples were determined by BCA Protein Analysis Kit (Thermo Fisher, TF268083). After being boiled for 5 min, the protein samples were electrophoresis by sodium dodecyl sulfate-polyacrylamide gel electrophoresis and then transferred to a polyvinylidene fluoride (PVDF) membrane, which was then blocked by incubation for 1 hour at room temperature in 5% milk in TBST. The blots were incubated overnight at 4°C with anti-USP25 antibody and anti-β-tubulin antibody. Then membrane was washed three times with TBST for 10 min each and incubated with goat anti-rabbit antibody or goat anti-mouse antibody for 40 min at room temperature. After incubation, the membrane was washed three times with TBST for 10 min each again.

### Co-immunoprecipitation

Co-immunoprecipitated cells were collected 48 h after transfection and washed once with normal saline. Lysis buffer were added to the six-well plate for 200μl-300μl/well (the normal saline had been precooled in advance. lysate: CO-IP lysate add protease inhibitor (100X) + Na3VO4 + DTT). After transfer to EP tube, cells were ultrasonic lysed at 15% power in the mode of 2s stop every 5s of ultrasound for a total of two minutes; After centrifugation for 15 min at 12 000 rpm and 4°C, 30μl supernatant was collected and boiled for 10 min with 30μl 2X loading as Input samples; the rest of the supernatant was transferred to another EP tube, followed by addition to 1μl/tube of IP antibody, incubation for 1-4 hours at 4°C, and addition to 25μl proteinA+G beads for overnight at 4°C, followed by centrifugation for 2min at 7500rpm and 4°C, suction with a suction pump to aspirate the supernatant, which can leave a little more liquid. Then the sample was added with 1ml IP high-salt wash, turned for 2min at 4°C, centrifugated for 2min at 7500rpm and 4°C. Beads were washed three times with high-salt wash buffer and three times with low-salt wash buffer. And at the last pass, IP samples were prepared aspirating the supernatant, remaining 30μl liquid or so, adding 30μl 2X loading, boiling in boiled water for 10min. Precipitates were fractionated using SDS-PAGE at appropriate concentrations. Perform experiments using the western blot as indicated.

### Ubiquitination assays

To analyze the ubiquitination of TRAF3 in HEK293T cells, plasmids encoding Tim3-red, USP25-myc, TRAF3-Flag and HA-ub-K48 were transfected into HEK293T cells for 48 hour and then treated with MG132 (20 μM), for 6 hours before harvesting.

eCells were lysed with immunoprecipitation lysis buffer (1% NP-40; Tris-HCl, 20 mM; NaCl, 150 mM; EDTA, 5mM; Na3VO4, 1mM; 0.5% and sodium deoxycholic acid and protease inhibitor cocktail, pH 7.5), and then whole-cell lysates were immunoprecipitated with Flag antibody (Sigma), followed by analysis of ubiquitination of TRAF3 with HA tag antibody. Precipitates were fractionated using SDS/PAGE at appropriate concentrations.

### Statistical analysis

The significance of the differences between groups was determined using a two-tailed Student’s t-test and a two-way ANOVA. For the mouse survival study, Kaplan-Meier survival curves were plotted and analyzed for statistically significance using GraphPad Prism 6.0. P value <0.05 was considered statistically significant.

## Notes

### Ethics statement

The animal study was approved by the Ethics Committee of Animal Experiments of the Beijing Institute of Basic Medical Sciences (IACUC-DWZX-2023-526). All efforts were made to minimize suffering.

---

**Fig S1.**
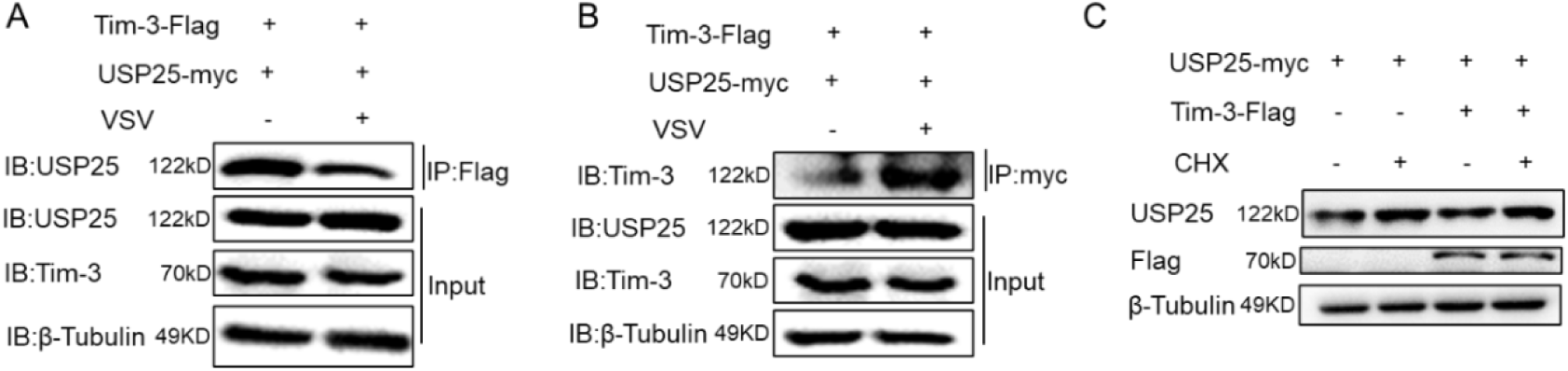
Tim-3 interacts with USP25. (A&B) HEK-293T cells were transfected with Tim-3-Flag plasmid and USP25-myc plasmid for 48 hours, then were left untreated or challenged by VSV for 12 hours, immunoprecipitated with Flag antibody (A) and with myc antibody (B), then detected by western blot for the indicated antibodies. (C) The HEK293T cells were transfected with plasmids encoding Tim-3-Flag and USP25-myc for 48 hours. Cells were treated with CHX (10 µM). Then cell lysis was used for immunoprecipitation and immunoblotting.

## References

[1] Han G, Chen G, Shen B, et al. Tim-3: an activation marker and activation limiter of innate immune cells[J]. Front Immunol, 2013, 4: 449. 10.3389/fimmu.2013.00449.

[2] Huang Y H, Zhu C, Kondo Y, et al. CEACAM1 regulates TIM-3-mediated tolerance and exhaustion[J]. Nature, 2015, 517(7534): 386–90. 10.1038/nature13848.

[3] Ju Y, Hou N, Meng J, et al. T cell immunoglobulin- and mucin-domain-containing molecule-3 (Tim-3) mediates natural killer cell suppression in chronic hepatitis B[J]. J Hepatol, 2010, 52(3): 322–9. 10.1016/j.jhep.2009.12.005.

[4] Sakuishi K, Jayaraman P, Behar S M, et al. Emerging Tim-3 functions in antimicrobial and tumor immunity[J]. Trends Immunol, 2011, 32(8): 345–9. 10.1016/j.it.2011.05.003.

[5] Ma H, Ren S, Meng Q, et al. Role of Tim-3 in COVID-19: a potential biomarker and therapeutic target[J]. Arch Virol, 2023, 168(8): 213. 10.1007/s00705-023-05842-2.

[6] Yang X, Jiang X, Chen G, et al. T cell Ig mucin-3 promotes homeostasis of sepsis by negatively regulating the TLR response[J]. J Immunol, 2013, 190(5): 2068–79. 10.4049/jimmunol.1202661.

[7] Jiang X, Zhou T, Xiao Y, et al. Tim-3 promotes tumor-promoting M2 macrophage polarization by binding to STAT1 and suppressing the STAT1-miR-155 signaling axis[J]. Oncoimmunology, 2016, 5(9): e1211219. 10.1080/2162402X.2016.1211219.

[8] Li G, Tang L, Hou C, et al. Peripheral Injection of Tim-3 Antibody Attenuates VSV Encephalitis by Enhancing MHC-I Presentation[J]. Front Immunol, 2021, 12: 667478. 10.3389/fimmu.2021.667478.

[9] Lee J, Su E W, Zhu C, et al. Phosphotyrosine-dependent coupling of Tim-3 to T-cell receptor signaling pathways[J]. Mol Cell Biol, 2011, 31(19): 3963–74. 10.1128/MCB.05297-11.

[10] Tian T, Li Z. Targeting Tim-3 in Cancer With Resistance to PD-1/PD-L1 Blockade[J]. Front Oncol, 2021, 11: 731175. 10.3389/fonc.2021.731175.

[11] Snyder N A, Silva G M. Deubiquitinating enzymes (DUBs): Regulation, homeostasis, and oxidative stress response[J]. J Biol Chem, 2021, 297(3): 101077. 10.1016/j.jbc.2021.101077.

[12] Meulmeester E, Kunze M, Hsiao H H, et al. Mechanism and consequences for paralog-specific sumoylation of ubiquitin-specific protease 25[J]. Mol Cell, 2008, 30(5): 610–9. 10.1016/j.molcel.2008.03.021.

[13] Nelson J K, Thin M Z, Evan T, et al. USP25 promotes pathological HIF-1-driven metabolic reprogramming and is a potential therapeutic target in pancreatic cancer[J]. Nat Commun, 2022, 13(1): 2070. 10.1038/s41467-022-29684-9.

[14] Zheng Q, Song B, Li G, et al. USP25 inhibition ameliorates Alzheimer’s pathology through the regulation of APP processing and Abeta generation[J]. J Clin Invest, 2022, 132(5). 10.1172/JCI152170.

[15] Zhu W, Zheng D, Wang D, et al. Emerging Roles of Ubiquitin-Specific Protease 25 in Diseases[J]. Front Cell Dev Biol, 2021, 9: 698751. 10.3389/fcell.2021.698751.

[16] Lin D, Zhang M, Zhang M X, et al. Induction of USP25 by viral infection promotes innate antiviral responses by mediating the stabilization of TRAF3 and TRAF6[J]. Proc Natl Acad Sci U S A, 2015, 112(36): 11324–9. 10.1073/pnas.1509968112.

[17] Ren Y, Zhao Y, Lin D, et al. The Type I Interferon-IRF7 Axis Mediates Transcriptional Expression of Usp25 Gene[J]. J Biol Chem, 2016, 291(25): 13206–15. 10.1074/jbc.M116.718080.

[18] Honda K, Takaoka A, Taniguchi T. Type I interferon [corrected] gene induction by the interferon regulatory factor family of transcription factors[J]. Immunity, 2006, 25(3): 349–60. 10.1016/j.immuni.2006.08.009.

[19] Seth R B, Sun L, Chen Z J. Antiviral innate immunity pathways[J]. Cell Res, 2006, 16(2): 141–7. 10.1038/sj.cr.7310019.

[20] Zhong B, Liu X, Wang X, et al. Ubiquitin-specific protease 25 regulates TLR4-dependent innate immune responses through deubiquitination of the adaptor protein TRAF3[J]. Sci Signal, 2013, 6(275): ra35. 10.1126/scisignal.2003708.

[21] Li J, Wang J, Pan T, et al. USP25 deficiency promotes T cell dysfunction and transplant acceptance via mitochondrial dynamics[J]. Int Immunopharmacol, 2023, 117: 109917. 10.1016/j.intimp.2023.109917.

[22] Teo Q W, Wong H H, Heunis T, et al. Usp25-Erlin1/2 activity limits cholesterol flux to restrict virus infection[J]. Dev Cell, 2023, 58(22): 2495–2509 e6. 10.1016/j.devcel.2023.08.013.

[23] Zhong B, Liu X, Wang X, et al. Negative regulation of IL-17-mediated signaling and inflammation by the ubiquitin-specific protease USP25[J]. Nat Immunol, 2012, 13(11): 1110–7. 10.1038/ni.2427.

[24] Gersch M, Wagstaff J L, Toms A V, et al. Distinct USP25 and USP28 Oligomerization States Regulate Deubiquitinating Activity[J]. Mol Cell, 2019, 74(3): 436–451 e7. 10.1016/j.molcel.2019.02.030.

[25] Sauer F, Klemm T, Kollampally R B, et al. Differential Oligomerization of the Deubiquitinases USP25 and USP28 Regulates Their Activities[J]. Mol Cell, 2019, 74(3): 421–435 e10. 10.1016/j.molcel.2019.02.029.

[26] Dou S, Li G, Li G, et al. Ubiquitination and degradation of NF90 by Tim-3 inhibits antiviral innate immunity[J]. Elife, 2021, 10. 10.7554/eLife.66501.

[27] Shi Q, Li G, Dou S, et al. Negative Regulation of RIG-I by Tim-3 Promotes H1N1 Infection[J]. Immunol Invest, 2023, 52(1): 1–19. 10.1080/08820139.2022.2113407.

[28] Van De Weyer P S, Muehlfeit M, Klose C, et al. A highly conserved tyrosine of Tim-3 is phosphorylated upon stimulation by its ligand galectin-9[J]. Biochem Biophys Res Commun, 2006, 351(2): 571–6. 10.1016/j.bbrc.2006.10.079.

[29] Gao P, Ma X, Yuan M, et al. E3 ligase Nedd4l promotes antiviral innate immunity by catalyzing K29-linked cysteine ubiquitination of TRAF3[J]. Nat Commun, 2021, 12(1): 1194. 10.1038/s41467-021-21456-1.

[30] Tang J C, Li Y, Wang Y L, et al. TRAF5 splicing variants associate with TRAF3 and RIP1 in NF-kappaB and type I IFN signaling in large yellow croaker Larimichthys crocea[J]. Fish Shellfish Immunol, 2022, 130: 418–427. 10.1016/j.fsi.2022.09.042.

[31] Tseng P H, Matsuzawa A, Zhang W, et al. Different modes of ubiquitination of the adaptor TRAF3 selectively activate the expression of type I interferons and proinflammatory cytokines[J]. Nat Immunol, 2010, 11(1): 70–5. 10.1038/ni.1819.

[32] Fernandez-Sesma A. The influenza virus NS1 protein: inhibitor of innate and adaptive immunity[J]. Infect Disord Drug Targets, 2007, 7(4): 336–43. 10.2174/187152607783018754.

[33] Lin C Y, Shih M C, Chang H C, et al. Influenza a virus NS1 resembles a TRAF3-interacting motif to target the RNA sensing-TRAF3-type I IFN axis and impair antiviral innate immunity[J]. J Biomed Sci, 2021, 28(1): 66. 10.1186/s12929-021-00764-0.

